# Subtilase activity in the intrusive cells mediates haustorium maturation in parasitic plants

**DOI:** 10.1101/2020.03.30.015149

**Authors:** Satoshi Ogawa, Takanori Wakatake, Thomas Spallek, Juliane K. Ishida, Ryosuke Sano, Tetsuya Kurata, Taku Demura, Satoko Yoshida, Yasunori Ichihashi, Andreas Schaller, Ken Shirasu

## Abstract

Parasitic plants that infect crops are devastating to agriculture throughout the world. They develop a unique inducible organ called the haustorium, which connects the vascular systems of the parasite and host to establish a flow of water and nutrients. Upon contact with the host, the haustorial epidermal cells at the interface with the host differentiate into specific cells called intrusive cells that grow endophytically towards the host vasculature. Then, some of the intrusive cells re-differentiate to form a xylem bridge that connects the vasculatures of the parasite and host. Despite the prominent role of intrusive cells in host infection, the molecular mechanisms mediating parasitism in the intrusive cells are unknown. In this study, we investigated differential gene expression in the intrusive cells of the facultative parasite *Phtheirospermum japonicum* in the family Orobanchaceae by RNA-Sequencing of laser-microdissected haustoria. We then used promoter analyses to identify genes that are specifically induced in intrusive cells, and used promoter fusions with genes encoding fluorescent proteins to develop intrusive cell-specific markers. Four of the intrusive cell-specific genes encode subtilisin-like serine proteases (SBTs), whose biological functions in parasitic plants are unknown. Expression of an SBT inhibitor in the intrusive cells inhibited their development, inhibited the development of the xylem bridge, and reduced auxin response levels near the site where the xylem bridge normally develops. Therefore, we propose that subtilase activity plays an important role in haustorium development in this parasitic plant.

**One sentence summary:** Tissue-specific analysis showed that the subtilases specifically expressed in intrusive cells regulate auxin-mediated host-parasite connections in the parasitic plant *Phtheirospermum japonicum*.

---

There are about 4500 species of parasitic plants; they are widespread, and those that infect crops are serious threats to agriculture (Yoshida *et al*., 2016; Clarke *et al*., 2019). In particular, members of the family Orobanchaceae, such as *Striga* spp. and *Orobanche* spp., are destructive root parasitic plants that invade major crops including rice, sorghum, and maize, often in resource-poor societies, and cause annual economic losses of over 1 billion U.S. dollars (Parker 2009, Runo and Kuria 2018). Parasitic Orobanchaceae plants produce large numbers of tiny seeds that are widely spread by wind, water, and people. To germinate, these seeds require host-derived stimulants such as strigolactones, which are a class of phytohormones (Yoneyama *et al*., 2010). These seeds can survive for decades in soil without germination, and thus it is difficult to eliminate parasitic plants from agricultural fields (Scholes and Press, 2008; Spallek *et al*., 2013; Gobena *et al*., 2017).

Parasitic plants develop a unique inducible organ called the haustorium that is used for invasion of the host plants. The haustorium connects the vasculature of the parasite with that of the host to establish a flow of nutritients and water from the host to the parasite (Yoshida *et al*., 2016; Clarke *et al*., 2019). Upon recognition of host-derived haustorium-inducing factors (Lynn and Chang, 1990), the parasite initiates organogenesis by activating cell division and cell expansion. In Orobanchaceae parasites, once the haustorium approaches the host, the epidermal cells in proximity to the host cells differentiate into intrusive cells, which have highly elongated shapes and function by intruding into the host (Musselman and Dickison, 1975). Once intrusive cells reach the host vasculature, some of the intrusive cells differentiate into xylem vessels, and subsequently formation of a xylem bridge (XB) between the parasite and host vasculature systems is initiated (Musselman and Dickison, 1975; Cui *et al*., 2016; Wakatake *et al*., 2018).

Despite many studies aimed at analyzing the transcriptional changes that occur during haustorium development and host infection in various species of parasitic plants (Ranjan *et al*., 2014; Yang *et al*., 2015; Zhang et al., 2015; Ichihashi *et al*., 2015, Sun *et al*., 2018; Yoshida *et al*., 2019), there have been few functional studies of these haustorium-specific genes. To explore the molecular mechanisms of parasitism, including haustorium organogenesis, we established a model parasitic plant system using *Phtheirospermum japonicum*, a facultative parasitic plant in the Orobanchaceae (Ishida *et al*., 2016; Spallek *et al*., 2017). *P. japonicum* is a self-fertilizing plant with a diploid genome, allowing forward genetics studies (Cui *et al*., 2016). In addition, an efficient root transformation system by *Agrobacterium rhizogenes*-mediated hairy root formation has been established, making functional studies of haustorial genes feasible (Ishida *et al*., 2011). To identify genes important for parasitism, we previously performed transcriptome analyses using rice-infecting *P. japonicum* and identified genes strongly expressed during the parasitic stage (Ishida *et al*., 2016). Among these was the auxin biosynthetic gene *YUCCA3*, which contributes to auxin biosynthesis in the haustorium. This auxin undergoes intercellular transportation and leads to the differentiation of tracheary elements, resulting in the formation of the XB that connects the parasite with the host (Ishida *et al*., 2016, Wakatake *et al*., 2018; Wakatake *et al*., 2020).

In this study, we identified differentially expressed genes in *P. japonicum* intrusive cells by using a laser microdissection method (LMD) combined with transcriptome analysis. We then used temporal and spatial promoter analyses to establish intrusive cell-specific gene markers. Among the upregulated genes, we focused on four genes encoding subtilisin-like serine proteases (subtilases; SBTs) that were exclusively expressed in the intrusive cells. We found that expression of an SBT inhibitor protein in the intrusive cells inhibited the maturation of the haustorium. Thus, our findings provide molecular insight about how parasitic plants develop their haustoria via SBTs.

## RESULTS

### Genes specifically expressed in the intrusive cells

Intrusive cells only form at the interphase between parasite and a susceptible host and thus likely participate in the invasion into host tissues and the molecular dialogue between parasite and host (Goyet *et al*., 2019). Despite the distinctive nature of intrusive cells, they have not been studied functionally and in detail yet. To seek molecular markers of their function in *P. japonicum*, we performed LMD coupled with tissue-specific transcriptome analysis. We used rice (*Oryza sativa* cv. Koshihikari) as the host plant because *P. japonicum* forms haustoria with relatively more intrusive cells on rice than on *Arabidopsis* roots (Fig. 1A, B). We separated the intrusive regions from other parts of the haustoria using LMD with cryosectioned haustoria (Fig. 1C, D), and obtained transcriptome profiles using the Illumina MiSeq system. After filtering out rice-derived sequences, the reads were mapped onto the draft genome of *P. japoncium* (Conn *et al*., 2015) and gene expression values were obtained. Whole transcriptome data are listed in Supplemental Data Set S1. We detected a total of 3079 differentially expressed genes between the intrusive cell region and the remainder of the haustorium (Supplemental Data Set S2). Subsequent Gene Ontology (GO) analysis revealed that nine GO terms, including “cell wall”, “response to biotic stimulus”, “transporter activity”, and “metabolic process” were enriched in both regions, whereas 15 GO terms, including “lipid metabolic process” and “carbohydrate metabolic process” were enriched specifically in other parts of the haustorium (Supplemental Tables S1 and S2). Only one term, “response to stress” was enriched in intrusive cells but not in other parts of the haustorium (Supplemental Table S1).

**Figure 1.**
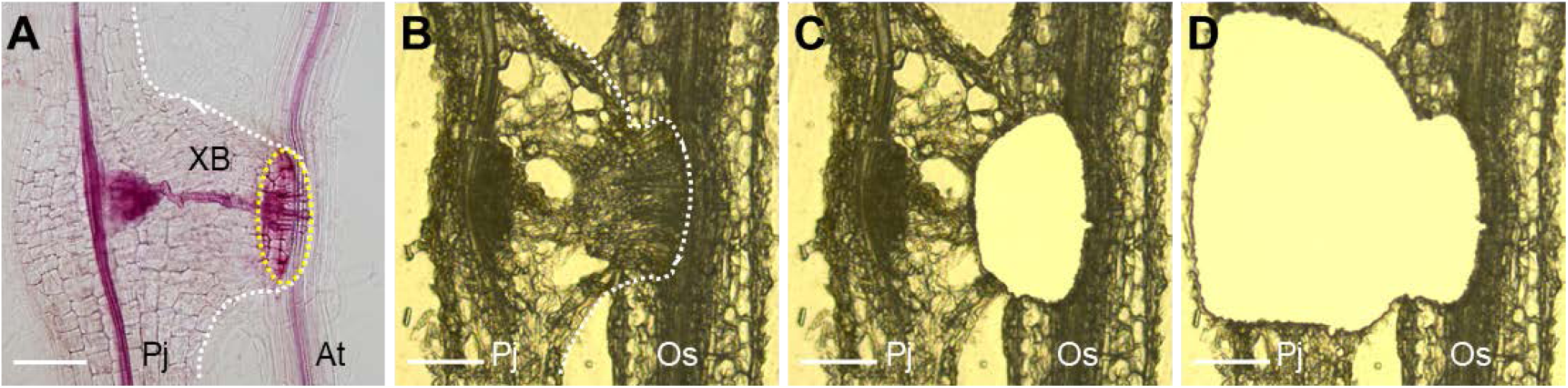
Laser microdissection of the haustorium in *P. japonicum*. (A) Safranin-O-stained A safranin-O-stained haustorium at 3 dpi. Within the haustorium, *P. japonicum* establishes a vascular connection with the *Arabidopsis* root, called the xylem bridge. The dashed white line outlines the haustorium. Intrusive cells located at the interface with the host (dashed yellow circle). (B-D) Sample preparation for tissue-specific transcriptome analysis of a *P. japonicum* haustorium infecting a rice root. Example of a cryosectioned haustorium before laser microdissection (B), after dissecting the intrusive region (C), and after dissecting the other part of the haustorium (D). Pj, *P. japonicum* root; At, *Arabidopsis thaliana* root; XB, xylem bridge; Os, *Oryza satiνa* root. Bar = 100 µm.

Next, we aimed to identify marker genes for intrusive cells as tools to investigate this cell type further. We selected three candidates among the differentially expressed genes that showed strong and specific expression in the intrusive cells: a homolog of *Haesa-like1* (*HSL1*) that we named *Intrusive Cell-Specific Leucine-rich repeat receptor-like kinase1* (*ICSL1*), *Germin-Like Protein1* (*GLP1*), and *Constitutive Disease Resistance1* (*CDR1*). These genes encode a leucine-rich repeat receptor-like kinase (LRR-RLK), a germin-like protein, and an aspartic protease, respectively (Xia *et al*., 2004; Ham *et al*., 2012; Qian *et al*., 2018). To test whether these genes show specific expression in intrusive cells, we made constructs containing each gene promoter linked to the sequence encoding a nuclear-localized fluorescent protein (3xVenus-NLS). We used the constructs to transform *P. japoncium* and analyzed the Venus fluorescence in *P. japoncium* haustoria formed after infection of *Arabidopsis thaliana* roots. For all constructs, fluorescence was detected specifically in the intrusive cells at 2 days post-infection (dpi) (Fig. 2A, C, E), and was stronger at 3 dpi (Fig. 2B, D, F). Intrusive cells are derived from epidermal cells, but an epidermis marker construct (*pAtPGP4::3xVenus-NLS*) is not expressed in the intrusive region (Wakatake *et al*., 2018). To further verify that *ICSL1* expression is specific to intrusive cells, we used the *ICSL1* promoter to drive a fluorescent marker module that localizes to the plasma membrane (3xmCherry-SYP; Wakatake *et al*., 2018). In *P. japoncium* haustoria that were transformed with both *pAtPGP4::3xVenus-NLS* and *pICSL1::3xmCherry-SYP*, we found mutually exclusive expression patterns for the two constructs at 4 dpi (Supplemental Fig. S1), with only *pICSL1::3xmCherry-SYP* expression in the intrusive cells. Based on these analyses, we defined that *ICSL1* as a reliable intrusive cell marker for further analyses.

**Figure 2.**
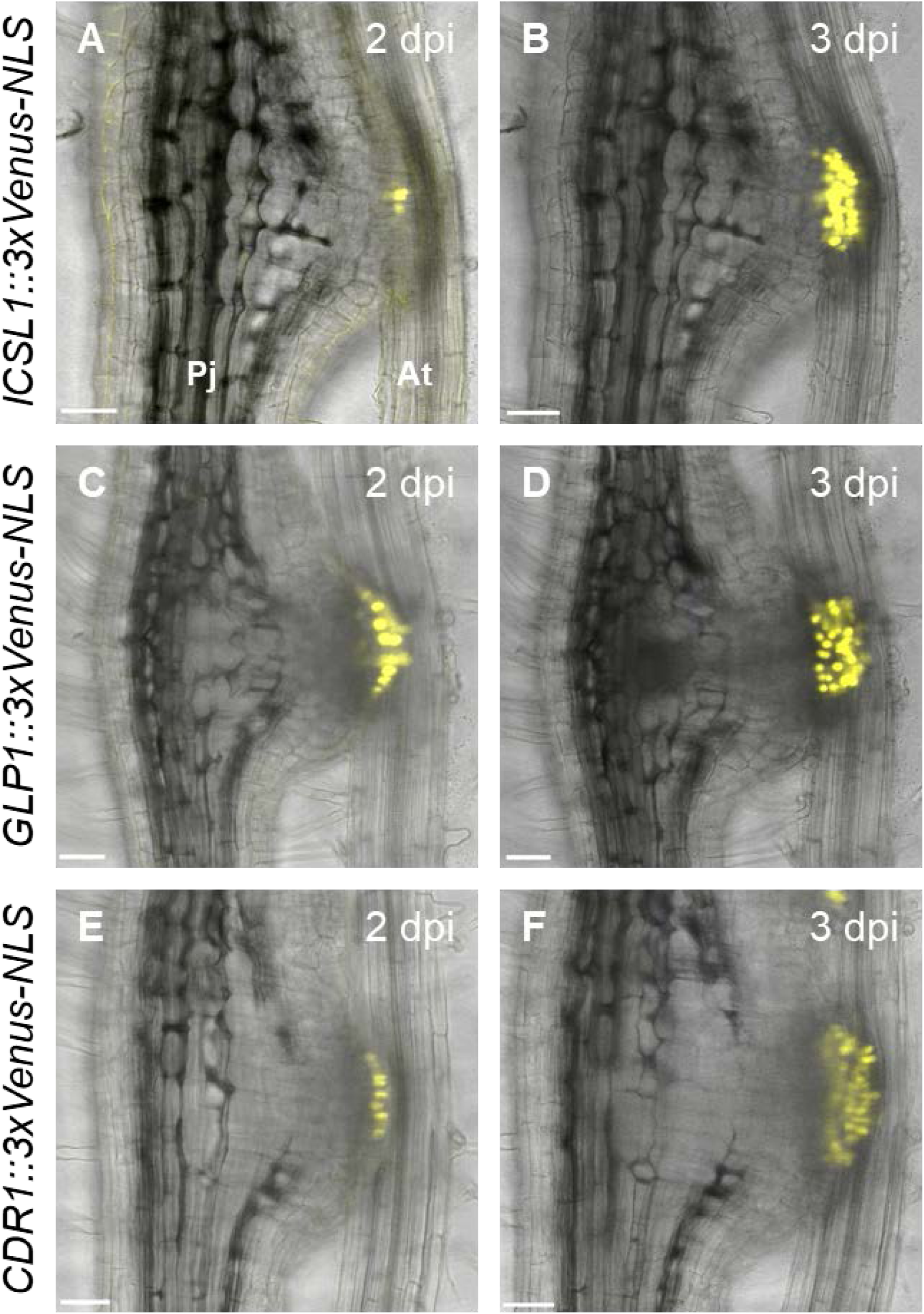
Expression dynamics of intrusive cell markers during haustorium development. Expression patterns of the *ICSL1* (A, B), *GLP1* (C, D), and *CDR1* (E, F) promoters driving a fluorescent marker gene during haustorium development at the indicated time points in *P. japonicum*. Bright-field and Venus fluorescent images were merged. Pj, *P. japonicum* root; At, *A. thaliana* root. Bar = 50 µm.

### Phylogeny and expression patterns of subtilases in *P. japonicum*

Among the genes that were expressed at higher levels in intrusive cells than in the remainder of the haustorium, we found five genes encoding subtilisin-like serine proteases (subtilases; SBTs). This was consistent with our previous report that *SBTs* are highly expressed during the parasitic stage in *P. japonicum* (Ishida *et al*., 2016). We therefore hypothesized that SBTs in intrusive cells may contribute to the host invasion process. To classify the *SBT* genes expressed in intrusive cells, we first identified all SBTs in the *P. japonicum* genome (Conn *et al*., 2015) on basis of their Asp-His-Ser catalytic triad and their peptidase S8 family domain (Smith *et al*., 1966; Wright *et al*., 1969). As a result, 97 putative SBTs met these criteria (Fig. 3). A phylogenetic analysis revealed that the five SBTs upregulated in intrusive cells all belong to Group 1 (Taylor and Qiu, 2017; Reichardt *et al*., 2018) (Fig. 3), which contains many SBTs involved in biotic interactions. These genes were thus designated as *SBT1.1.1, SBT1.2.3, SBT1.5.2, SBT1.7.2*, and *SBT1.7.3*. We also found that the many of the 97 *SBT* genes in *P. japonicum* were induced in the haustorium at 3 days post-infection (dpi) or later (Fig. 3), indicating that these *SBTs* were activated after attachment to the host.

**Figure 3.**
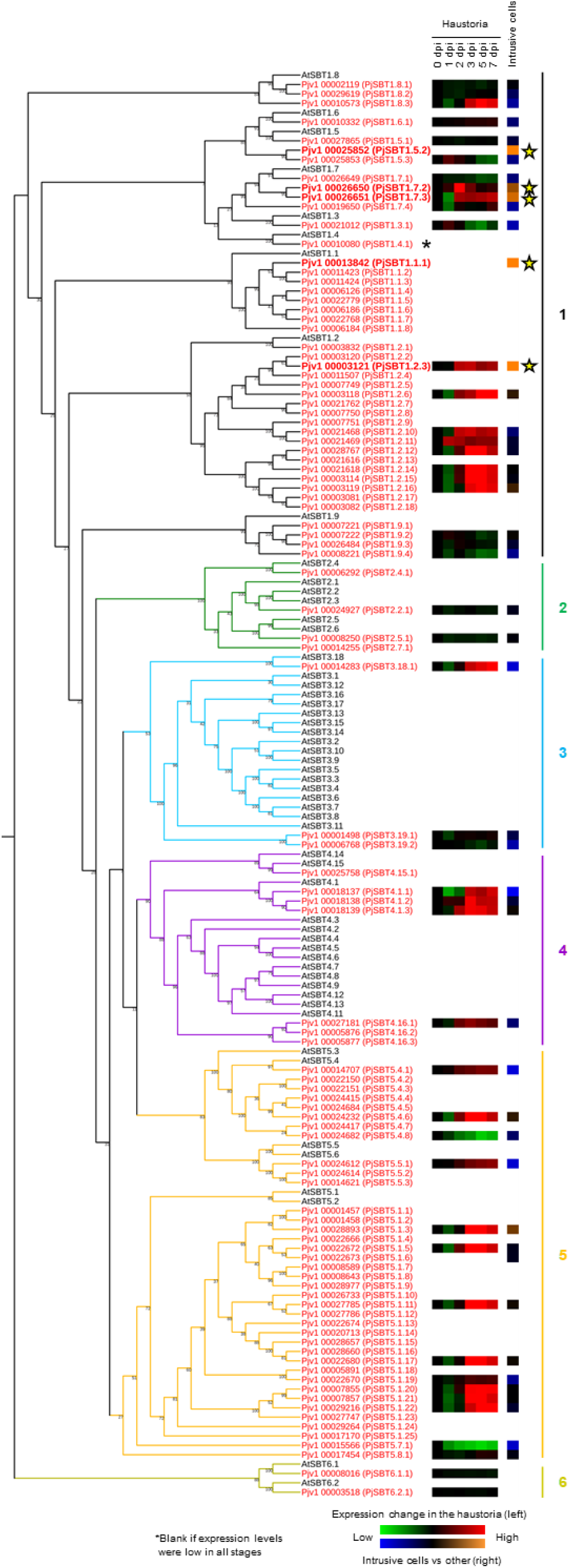
Phylogeny of the subtilases (SBTs) in *P. japonicum* and *Arabidopsis*. The 97 SBTs in *P. japonicum* are shown in red and the 55 SBTs in *Arabidopsis* are shown in black. According to Rautengarten *et al*. (2005), the SBTs are categorized into 6 groups. The group 6 SBTs represent the outgroup. The green/red squares indicate the *SBT* gene expression levels at different time points in the haustorium of *P. japonicum*. The blue/orange squares indicate the *SBT* gene expression levels in the intrusive cells relative to their expression in other haustorial parts. Stars indicate the *SBT*s with higher expression in the intrusive cells than in the other haustorial parts.

### Subtilases specifically expressed in intrusive cells

To confirm the expression patterns of the five *SBTs* up-regulated in intrusive cells, we made constructs containing each gene promoter linked to the 3xVenus-NLS module, transformed *P. japoncium* with the constructs, and analyzed Venus fluorescence in *P. japoncium* haustoria after infection of *A. thaliana* roots with a confocal microscope. The Venus signal driven by the putative *SBT1.5.2* promoter was not detected at selected time points. We thus focused on the remaining four *SBTs* in further analyses. An alignment of their protein products is shown in Supplemental Fig. S2. The promoters of *SBT1.1.1, SBT1.2.3*, and *SBT1.7.3* were sufficient to drive detectable Venus expression in the intrusive cells at 3 to 7 dpi (Fig. 4A). Expression of *SBT1.7.2* was more transient, with weaker signal at 7 dpi as compared to 3 and 5 dpi. For the *SBT1.7.3* promoter, signals were detected also in vascular cells in the meristematic region (Fig. 4A). We used quantitative reverse transcription PCR (RT-qPCR) to analyze induction of the four *SBTs* in whole haustoria, and found that the levels of induction at 3 and 7 dpi were consistent with the results from the Venus fluorescence analysis (Fig. 4B). We also analyzed expression of a 3xVenus-NLS construct driven by the *SBT1.7.1* gene, which is phylogenetically close to *SBT1.7.2* and *SBT1.7.3* (Fig. 3). Florescence from this construct was observed in the epidermal cells but not the intrusive cells (Supplemental Fig. S3). Since the intrusive cells are uniquely found in parasitic plants, *SBT1.1.1, SBT1.2.3, SBT1.7.2*, and *SBT1.7.3* expression in this cell-type suggest that these *SBTs* function in parasitism.

**Figure 4.**
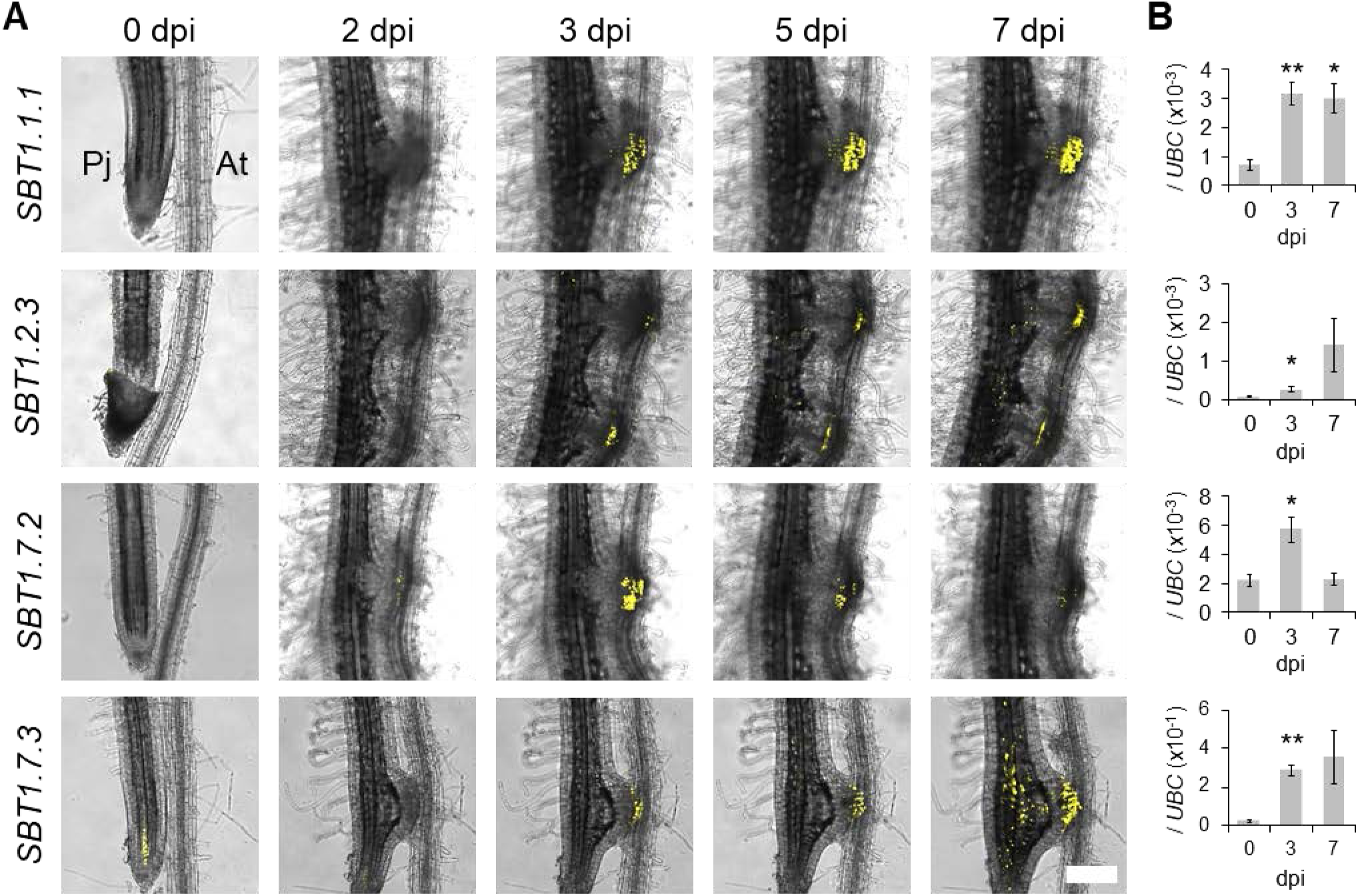
Expression dynamics of the SBTs during haustorium development. (A) Expression patterns of *SBT* promoters in *P. japonicum* during haustorium development at the indicated time points. Bright-field and Venus fluorescent images were merged. Pj, *P. japonicum* root; At, *A. thaliana* root. Bar = 200 µm. (B) The relative expression level of each *SBT* at 0, 3, and 7 dpi in isolated haustoria. “0 dpi” values represent the expression of the marker genes in the root elongation zones of non-infecting *P. japonicum* roots. Representative data are shown (mean ± SE of 4 technical replicates). We used *PjUBC* as a reference gene. The experiments were performed three times with similar results. Asterisks indicate statistical significance (*P < 0.05, **P < 0.01).

### Subtilases play important roles in development of the host-parasite connection via auxin signaling

The four *SBT* genes discussed above may be functionally redundant, and silencing multiple genes in *P. japonicum* is challenging due to the lack of a transgenerational transformation method. Therefore, we used an SBT inhibitor protein to analyze the functions of the intrusive cell-specific SBTs. For this purpose, we chose <u>E</u>xtracellular <u>p</u>roteinase inhibitor 10 (Epi10) from *Phytophthora infestans*. Epi10 carries an atypical Kazal domain and inhibits subtilases but does not inhibit the other major serine proteases, trypsin and chymotrypsin (Tian and Kamoun, 2005; Tian *et al*., 2005). The tissue-specific inhibition of SBTs has previously been accomplished by expressing *Epi10* under a tissue-specific promoter (Schardon *et al*., 2016). To specifically inhibit the SBTs expressed in developing haustoria, we used the promoter sequences of *SBT1.1.1* and *SBT1.2.3* to drive expression of the *Epi10* coding region. We compared the development of haustoria in *P. japonicum* roots transformed with these constructs with the development of haustoria in control roots transformed with an empty vector. We found that hairy roots transformed with the *Epi10* constructs showed reduced xylem bridge (XB) formation in the haustoria at 5 dpi after infection of *Arabidopsis* roots when compared with control hairy roots (Fig. 5A–C, Supplemental Fig. S4).

**Figure 5.**
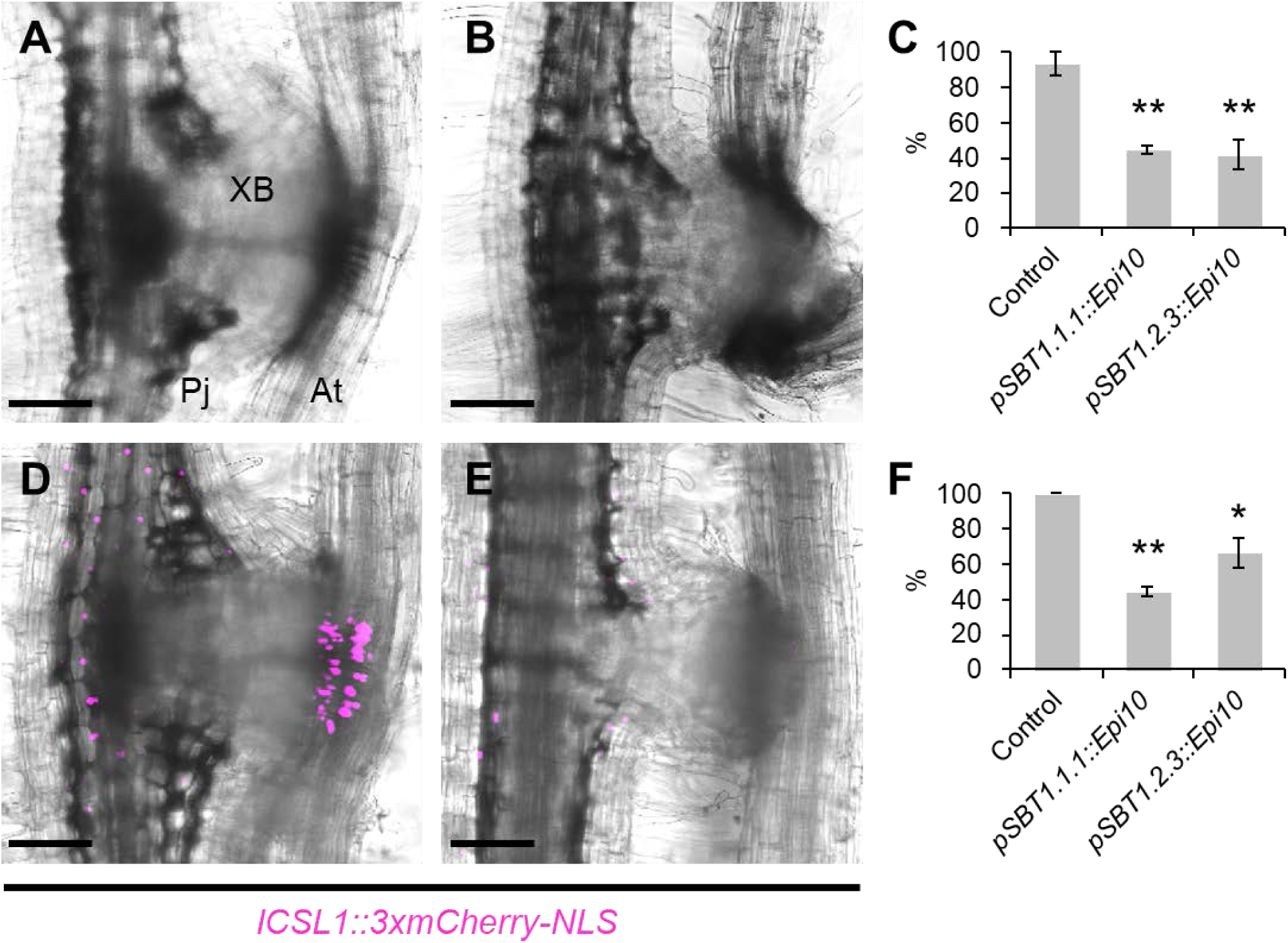
Effect of the SBT inhibitor Epi10 on haustorial development. (A) Representative image of a control haustorium that formed a xylem bridge (XB) at 5 dpi. (B) Representative image of a haustorium expressing the SBT inhibitor Epi10 that did not form an XB at 5 dpi. (C) Ratios of haustoria that formed XBs in *P. japonicum* plants transformed with the empty vector (Control) or with the *Epi10* gene driven by the *SBT1.1.1* or *SBT1.2.3* promoters (mean ± SE of 3 replicates, n = 4–7). (D, E) Representative images of haustoria that did (D) and did not (E) form intrusive cells at 5 dpi. The *ICSL1* promoter was used as an intrusive cell marker. The bright-field and mCherry fluorescent images are merged. (F) Ratios of haustoria that formed intrusive cells in *P. japonicum* plants transformed with the empty vector (Control) or with the *Epi10* gene driven by the *SBT1.1.1* or *SBT1.2.3* promoters (mean ± SE of 3 replicates, n = 4–7). Asterisks indicates statistical significance (*P < 0.1, **P < 0.01). Pj, *P. japonicum*; At, *A. thaliana*; XB, xylem bridge. Bar = 100 µm.

Next, we investigated whether the *Epi10*-transformed hairy roots would show other developmental abnormalities. We were particularly interested in the effects on intrusive cells, given the specific expression of *SBT1.1.1* and *SBT1.2.3* in intrusive cells. Therefore, we monitored expression of the intrusive cell marker *ICSL1* (Fig. 2A, B) in the *Epi10*-expressing haustoria. To accomplish this, we transformed *P. japonicum* roots with each of the *Epi10* constructs and with a construct encoding the mCherry fluorescent protein with a nuclear localization signal (3xmCherry-NLS) driven by the *ICSL1* promoter. In control roots transformed with *pICSL1::3xmCherry-NLS* but not with *Epi10*, all haustoria showed specific mCherry fluorescence in the intrusive cells. In contrast, less than 45% of the *pSBT1.1.1::Epi10* haustoria, and approximately 67% of the *pSBT1.2.3::Epi10* haustoria showed mCherry fluorescence at 5 dpi (Fig. 5D–F). These results suggest that the intrusive cell-specific SBT activities promote the maturation of haustoria by regulating the development of intrusive cells and the subsequent XB formation. Lack of intrusive cell identity may affect auxin distribution (Ishida *et al*., 2016, Wakatake *et al*., 2020). Therefore, we investigated whether *Epi10* expression alters auxin signaling within the haustorium by using the 3xmCherry-NLS module controlled by the synthetic, auxin-responsive *DR5* promoter (Ulmasov *et al*., 1995). Most of the auxin signaling in the central region of the haustoria, but not around the xylem plate, was diminished by Epi10 (Fig. 6). Taken together, our results suggest that the SBT activities regulate auxin-dependent maturation of *P. japonicum* haustoria.

**Figure 6.**
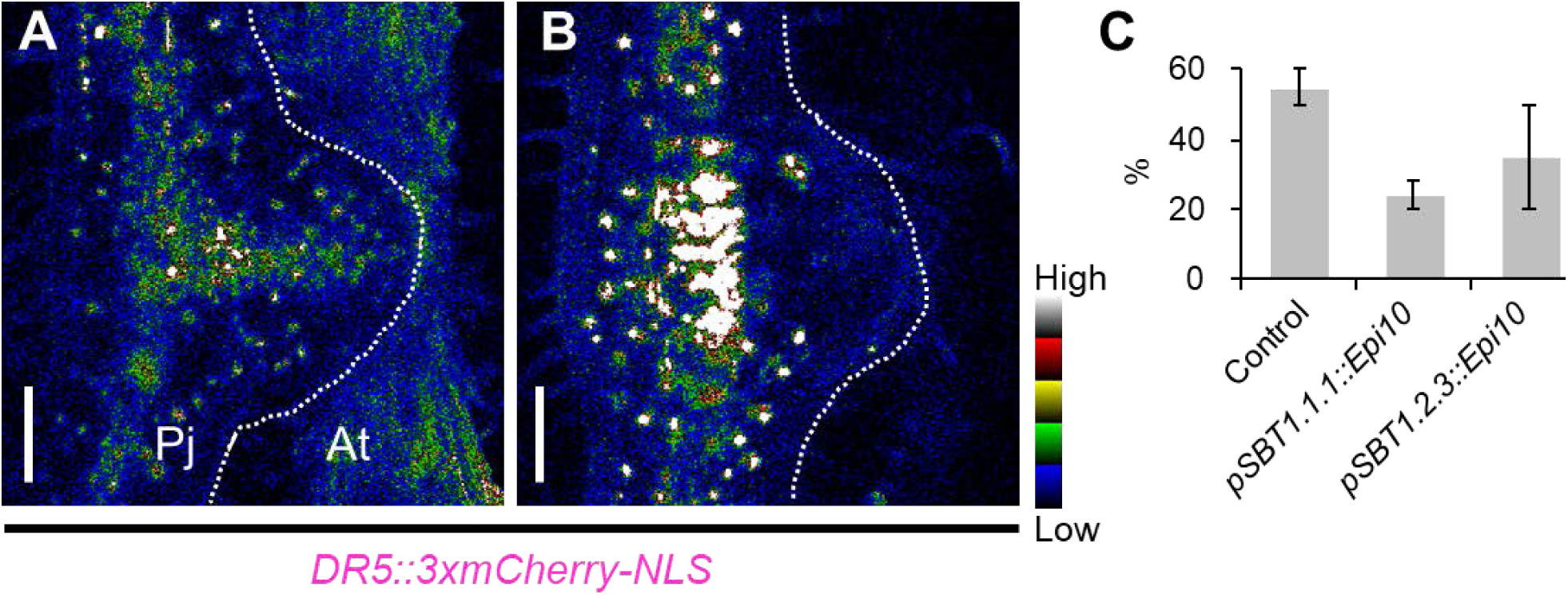
Effect of the SBT inhibitor on auxin signaling. (A, B) Representative images of haustoria in which auxin signaling was observed (A) and not observed (B) around the xylem bridge at 5 dpi. The *DR5* promoter was used as an auxin signaling marker. The mCherry fluorescence intensity is depicted in a 5 ramps spectrum. The broken white line indicates the edge of the haustorium. (C) Ratios of haustoria in which auxin signaling was observed around the xylem bridge in *P. japonicum* plants transformed with the empty vector (Control) or with the *Epi10* gene driven by the *SBT1.1.1* or *SBT1.2.3* promoters (mean ± SE of 2 replicates, n = 2–7). Pj, *P. japonicum*; At, *A. thaliana*; XB, xylem bridge. Bar = 100 µm.

## DISCUSSION

We used *P. japonicum* as a model parasitic plant to elucidate the molecular mechanisms that regulate parasitic functions in the intrusive cells of the haustoria. By using tissue-specific RNA-seq analysis coupled with LMD, we identified a number of genes that are up-regulated in intrusive cells (Supplemental Data Set S1). A previous study used the LMD method to reveal genes that are specifically expressed at the host-parasite interface, which includes the intrusive cells, in the facultative hemiparasite *Tryphisaria versicolor* infecting *Zea mays* or *Medicago truncatula* (Honaas *et al*., 2013). In that study, the GO term “transcription factor activity” was overrepresented, while the term “transporter activity” was underrepresented at the host-parasite interface. In contrast, we found that the term “transporter activity” was enriched in the intrusive cells in *P. japonicum* (Supplemental Table S1). Although a direct comparison of the two experiments is difficult, the results may indicate that *T. versicolor* and *P. japonicum* employ different strategies to invade their particular hosts. Enrichment of “transporter activity” in intrusive cells also suggests that these cells may have a function in material transfer, which would be consistent with their position at the interface between host vasculature and parasite haustorium. We found that several GO terms, such as “lipid metabolic process” and “carbohydrate metabolic process”, are strongly enriched in the rest of the haustorium but not in the intrusive cells (Supplemental Tables S1 and S2). Yoshida *et al*. (2019) showed that genes categorized under the GO terms “protein metabolic process”, “carbohydrate metabolic process”, and “catabolic process” are upregulated in rice-infecting *Striga hermonthica* at 7 dpi, when the host-parasite connection in the haustorium is established. This result indicates that metabolically demanding processes such as morphology are activated in the haustoria in the family Orobanchaceae.

Based on our intrusive cell-specific transcriptome, we established that the three *P. japonicum* genes *ICSL1, GLP1*, and *CDR1* showed strong and specific expression in the intrusive cells and could be used as molecular markers for these cells (Fig. 2). ICSL1 is homologous to the *Arabidopsis* HSL1 receptor, which localizes to the plasma membrane and recognizes peptide hormones (Torii, 2004; Macho and Zipfel, 2014; Shinohara *et al*., 2016). The phylogenetically closest *Arabidopsis* ICSL1 homolog, however, is AtRLP52, a receptor-like kinase associated with disease resistance and an unknown ligand (Ramonell *et al*., 2005; Ellendorff *et al*., 2008) (Supplemental Fig. S5). Thus, it is possible that ICSL1 also recognizes peptide hormones. Further experiments are required to identify the unknown ICSL1 ligand and to determine if it originates from the parasite or the host. The second marker gene encodes GLP1, which belongs to a GLP superfamily, which consists of structurally diverse plant glycoproteins including enzymes such as oxalate oxidases and superoxide dismutases (Rietz *et al*., 2012; Sakamoto *et al*., 2015). Phylogenetic analysis revealed that GLP1 in *P. japonicum* is closely related to *Arabidopsis* GLP1 and GLP3, which lack oxalate oxidase activity, and to GhABP19 in *Gossypium hirsutum*, a superoxide dismutase potentially regulating redox status (Pei *et al*., 2019) (Supplemental Fig. S6). Interestingly, the only other gene with experimentally confirmed expression in intrusive cells encodes a peroxidase in *S. hermonthica* (Yoshida *et al*., 2019). Also, chemically inhibiting peroxidase activity, and thus altering the redox homeostasis, reduces haustorium formation in *Striga* spp. and *Triphysaria* (Wada *et al*., 2019; Wang *et al*., 2019). The expression of a superoxide dismutase in *P. japonicum* intrusive cells further supports a role for redox regulating enzymes in haustorium development. The third intrusive cell-specific gene that we identified encodes CDR1, which belongs to a family of aspartic proteases. The *P. japonicum* CDR1 is a homolog of aspartic proteases that regulate disease resistance signaling in *Arabidopsis* (Xia *et al*., 2004) (Supplemental Fig. S7). It is currently not known if CDR1 regulates defense responses in *P. japonicum;* however, the expression of defense-related genes in *P. japonicum* haustoria was seen in a previous microarray study (Ishida *et al*., 2016).

*ICSL1, GLP1*, and *CDR1* show the same spatio-temporal expression pattern, with detectable expression beginning at 2 dpi at the interface with the host (Fig. 2A, C, E). This is the time point when expression of an epidermis marker ceases in the same region (Wakatake *et al*., 2018). Thus, the developmental switch from epidermis to intrusive cell is likely to be activated around this time point. Considering the mutually exclusive expression patterns of the epidermis marker gene and the intrusive cell marker gene (Supplemental Fig. S1), we would expect that the transcriptional landscapes of these two cell types are substantially different. Specific expression of *SBT1.7.1* in the epidermal cells, but not in the intrusive cells support this idea further (Fig. S3). The intrusive cell-sepcific markers identified in this study were expressed uniformly in the entire intrusive region (Fig. 2). However, only a fraction of those cells differentiate into tracheary elements to be part of the XB (Wakatake *et al*., 2020). Thus it seems that there are different types of cells in the intrusive cell population. This is likely due to non-uniform auxin response in the intrusive region. Thus further detailed analyses are required to reveal mechanisms by which auxin responses are controlled in intrusive cells.

A transcriptome analysis of *P. japonicum* haustoria in our previous study revealed that 7 of the 10 genes with the highest, exclusive expression in the parasitic stage were *SBTs* (Ishida *et al*., 2016) (Supplemental Table S3). Many *SBT* genes are also upregulated in *Striga* spp. upon infection (Yoshida et al., 2019). SBTs are a widespread protein family existing in eubacteria, archaebacteria, eukaryotes, and viruses (Rawlings and Barrett, 1994; Schaller *et al*., 2018). In plants, SBTs are required for the maturation of plant peptide hormones, leading to phenotypic changes such as root elongation, abscission of floral organs, and embryonic cuticle integrity (Matsubayashi, 2014; Ghorbani *et al*., 2016; Schardon *et al*., 2016; Doll *et al*., 2020; Reichardt *et al*., 2020). Here, we identified four *SBTs* that are exclusively expressed in intrusive cells (Figs. 3 and 4), and they all belong to Group 1. Group 1 SBTs and Group 5 SBTs are highly expanded in parasitic plants compared with those in *Arabidopsis*, while Groups 3 and 4 are much smaller in *P. japonicum* than in *Arabidopsis* (Fig. 3, Supplemental Fig. S8). More than 40% of SBTs in the parasites *P. japonicum, Striga asiatica* and *S. hermonthica* belong to Group 1 SBTs. In addition, Group 1 SBTs also expanded in plants that undergo symbiosis with nitrogen-fixing bacteria. Group 1 SBTs also include many that are involved in plant defense (Taylor and Qiu, 2017; Reichardt *et al*., 2018). These findings indicate that Group 1 SBTs may have evolved for biotic interactions, including parasitism. The molecular functions and substrates of several Group 1 SBTs in non-parasitic plants have been investigated. For example, Phytaspase 2, a Group I SBT in tomato, cleaves and activates the peptide hormone PHYTOSULFOKINE (PSK), which induces stress-induced flower drop in tomato, in addition to its well-established growth regulatory and immune-modulating activities (Reichardt *et al*., 2020). Genes for PSK and its candidate receptor are present in the *P. japonicum* genome. Interestingly, expression of these two genes is upregulated in the haustoria but not in the intrusive cells. If SBT1.1.1 or SBT1.2.3 is involved in PSK precursor processing, the expression of these two proteins in two different cell types would suggest that they may facilitate the communication between haustorial tissues. A similar tissue-tissue dialogue mediated by SBTs was recently shown to operate during *Arabidopsis* seed development (Doll *et al*., 2020). In contrast, *Arabidopsis* SBT1.2 (alias SDD1), a homolog of *P. japonicum* SBT1.2.3, contributes to stomatal development (von Groll *et al*., 2002). The substrates of SDD1 have not been identified, but were suggested to also include plant peptide hormones.

We showed that SBT activity in intrusive cells contributes to haustorium development (Figs. 4-6, Supplemental Fig. S4). Intrusive cells *per se* were still formed in Epi10-transgenic hairy roots (Fig. 5B, E; Supplemental Fig. S4). Thus, we can hypothesize that SBTs contribute to differentiation of intrusive cells into xylem vessels, leading to XB formation. The specific expression of SBTs during the parasitic stage is shared between *P. japonicum* and *S. hermonthica*, suggesting that SBTs are important for parasitism in the family Orobanchaceae. Intrusive cell-specific SBTs have not yet been identified in *S. hermonthica.* However, an *S. hermonthica* SBT is expressed specifically in the haustorial hyaline body (Yoshida *et al*., 2019). The hyaline body consists of parenchymatic tissue in the central region of the haustorium and is characterized by dense, organelle-rich cytoplasm, abundant paramural deposits, and high metabolic activity (Visser *et al*., 1984). The hyaline body has not yet been identified in *P. japonicum* and it may be morphologically distinct from that in *S. hermonthica*. The further identification of cell-type-specific SBTs in haustoria may facilitate the identification and functional studies of the hyaline body in *P. japonicum*.

Our data suggest that expression of the *SBTs* may be initiated in cells that eventually become intrusive cells, then the SBT activities contribute to the maturation of the intrusive cells, where the marker gene *ICSL1* is expressed. After expression of intrusive cell-specific markers, intrusive cells may invade host tissue, intrusive cells reach the host vasculature, auxin is transported inward towards the root vasculature, and then the XB is formed (Wakatake *et al*., 2020). Importantly, treatment with haustorium-inducing factors induces organogenesis of the haustorium in *P. japonicum* without hosts, but the intrusive cells and XB are not formed in these haustoria (Ishida *et al*., 2016, Goyet *et al*., 2019). Identification of the unknown host-derived signals required for intrusive-cell specific SBT induction will provide insights into mechanisms of how Orobanchaceae parasites invade the host plants.

We also found that several *SBTs* were induced at the later stages of the infection both in *P. japonicum* and in *S. hermonthica* (Supplemental Fig. S8). The late expression of SBTs during the infection indicates that parasitic plants utilize SBTs also after attachment to the host, possibly in regulating parasitism. We focused our study on SBTs with a role in haustorium development that can be studied with transgenic *P. japonicum* hairy roots (Ishida *et al*., 2016; Wakatake *et al*., 2020). To address the role of SBTs in later stages of the infection would require the generation of stable transgenic plants. In addition, many SBT clades were found to be species-specific, suggesting that each parasite has recruited SBTs independently to promote parasitism. Since parasitic plants are able to transfer molecules such as phytohormones and microRNAs (Spallek *et al*., 2017; Shahid *et al*., 2018), it is possible that peptides processed by SBTs in the haustorium can be transported from the parasite into the host. Further analyses of peptides in infected hosts will be required to assess this hypothesis. In summary, our study showed that SBTs are required for haustorium development. Functional studies of other parasite SBTs and their targets will provide important insights into parasitism in future studies.

## MATERIALS AND METHODS

### Plant materials and growth conditions

*P. japonicum* (Thunb.) Kanitz and rice (*Oryza sativa* L. subspecies *japonica*, cv Koshihikari) seeds were handled as described previously (Yoshida and Shirasu, 2009; Ishida *et al*., 2011). For *in vitro* germination, *P. japonicum* seeds were surface sterilized with 10% commercial bleach solution (Kao, Tokyo, Japan) for 5 minutes, followed by 5 rinses with sterilized water. Seeds were then sown on solid half-strength MS medium (0.8% Bacto agar, 1% sucrose, pH 5.8). After stratification at 4°C in the dark overnight, plants were grown either vertically for infection assays or horizontally for transformation, at 25°C under long-day conditions (16-h light, 8-h dark). *Arabidopsis* (*Arabidopsis thaliana*, ecotype Col-0) seeds were surface sterilized with 5% commercial bleach solution for 5 minutes, followed by 5 rinses with sterilized water. Seeds were then sown on solid half-strength MS medium. After stratification at 4°C in the dark overnight, plants were grown vertically at 22°C under long-day conditions. Rice seeds were sterilized with 70% ethanol for 3 minutes, followed by incubation in a 50% commercial bleach solution for 20 minutes. After 5 rinses with sterilized water, seeds were sown on quarter strength Gamborg’s B5 medium (Sigma) with 0.7% agar (INA). Plates were kept vertically at 26°C under long-day conditions.

### Sample preparation for RNA-seq

Ten-day-old *P. japonicum* seedlings were transferred to quarter-strength Gamborg’s B5 medium (0.7% agar; INA) and grown vertically at 25°C under long-day conditions for 2 days. These seedlings and 7-day-old rice seedlings were transferred together to new quarter strength Gamborg’s B5 plates for infection at 25°C under long-day conditions. At 5 dpi, haustoria were excised and immediately soaked in chilled RNAlater (Sigma) and stored at 4°C. Samples were embedded in FSC 22 frozen section media (Leica biosystems) in self-made aluminum molds in an acetone bath at -75°C. Frozen blocks were sectioned to 20 µm thickness using a cryostat (Leica CM3050S) with adhesive seals at -30°C. Sections were transferred to room temperature and immediately air-dried. The intrusive regions and the other parts of the haustorium were dissected using a Leica LMD7000. Dissected tissues were collected in the lids of 0.5 mL microtubes filled with RNA extraction buffer. Approximately 20 haustoria were used for one biological replicate. Total RNAs were extracted using the Picopure RNA isolation kit (Arcturus) according to the manufacturer’s instructions. DNase I (Qiagen) was applied to the column during the procedure to digest genomic DNA. Elution buffer (11 µL) was used to elute the total RNA. The quality and quantity of the total RNA were assessed using a Bioanalyzer (Agilent Technologies) and the RNA 6000 pico kit.

### Whole-transcripts amplification and library preparation

The procedure for whole-transcript amplification was based on the Quartz-seq method (Sasagawa *et al*., 2013). Approximately 1 ng of total RNA was used as a starting material. Total RNA was denatured (70°C for 90 sec) and primed (35°C for 15 sec) followed by first-strand synthesis (35°C for 5 min; 45°C for 20 min; 70°C for 10 min) with reverse transcriptase and oligo-dT-containing RT primers, using the SuperScript III system (Life Technologies). Single stranded cDNA was purified using AMPure XP magnetic beads (Beckman Coulter). The remaining RT primers were digested on the beads with Exonuclease I (TAKARA) (37°C for 30 min; 80°C for 20 min). Subsequently, the poly-A-tailing reaction was performed with terminal transferase (Roche) (37°C for 50 sec; 65°C for 10 min) followed by second strand synthesis using a tagging primer with MightyAmp DNA polymerase (TAKARA) (98°C for 130 sec; 40°C for 1 min; 68°C for 5 min). PCR enrichment was performed using the enrichment primers and MightyAmp DNA polymerase (98°C for 10 sec; 65°C for 15 sec; 68°C for 5 min). The total number of PCR cycles was either 14 or 15, depending on the amount of total RNA input. The amplified cDNA was purified using DNA concentrator-5 (Zymo Research) according to the manufacturer’s instructions. The size distribution of the amplified cDNA was assessed using the Bioanalyzer with a High sensitivity DNA kit. The amount of cDNA in each sample was measured using the Qubit dsDNA HS assay kit (Thermo Fisher). Library preparation was performed using the Nextera XT kit (Illumina) according to the manufacturer’s instructions. The KAPA LA kit (Nippon Genetics) was used for library amplification after fragmentation. PCR cycles were adjusted to 8 or 9 depending on the amount of input cDNA. Libraries were pooled and sequenced with 3 runs on the MiSeq using the reagent kit V2 (Illumina).

### Bioinformatics analysis

The adapter sequences in the primers for library preparation and whole transcript amplification were trimmed and low-quality sequences were removed. Quality-filtered reads were mapped to Nipponbare-reference-IRGSP-1.0 pseudomolecules (Kawahara *et al*., 2013) using the CLC genomics workbench (ver. 8.0, Qiagen) with a threshold setting of 95% match. The remaining unmapped reads were considered as *P. japonicum*-derived sequences and mapped to the *P. japonicum* draft genome with a threshold setting of 90% match. Unique read counts obtained for each gene model were used for further analysis. Differential gene expression analysis was performed in R with the TCC package (Sun *et al*., 2013) (https://www.R-project.org/.). Gene ontology analysis was performed with GO seq using the results of the differential gene expression analysis (Young *et al*., 2010). We used the CLC Main Workbench (ver. 8.0.1, Qiagen) for identification of the putative SBTs in *P. japonicum*, vector design, and sequence analyses.

### Phylogenetic analysis

Phylogenetic analyses were performed using the CLC Genomics Workbench (ver. 8.0, Qiagen). Predicted amino acid sequences were trimmed using trimAL (Capella-Gutiérrez *et al*., 2009), followed by alignment. Based on the alignment, the phylogenetic tree was drawn using the maximum-likelihood method. For comparing the SBTs in *P. japonicum* and Arabidopsis, the reliability of the trees was tested by bootstrap analysis with 1000 resamplings. For comparing the SBTs in *P. japonicum, S. asiatica*, and *S. hermonthica*, the reliability of the trees was tested by bootstrap analysis with 100 resamplings. The figure was generated by iTOL (ver. 5) (https://itol.embl.de/)(Letunic and Bork, 2007).

### Cloning

Golden Gate cloning technology was used for cloning (Engler *et al*., 2014). All the *Bpi*I and *Bsa*I restriction sites within the cloned DNA sequences were mutated. The golden gate modules *3xVenus-NLS, 3xmCherry-SYP, pACT::3xmCherry-NLS*, and *pAtPGP4::3xVenus-NLS* were described previously (Ishida *et al*., 2016; Wakatake *et al*., 2018).

#### Vectors containing intrusive cell markers

The *PjICSL1* (2652 bp), *PjGLP1* (2634 bp), and *PjCDR1* (2496 bp) promoter regions were each PCR-amplified as two fragments from *P. japonicum* genomic DNA and cloned separately into pAGM1311. The fragments were then combined into the pICH41295 level 0 vector. The promoter sequences were next assembled into level 1 vectors together with the fluorescent protein module and the 3’UTR and *HSP* terminator module. *pICSL1::3xmCherry-SYP* was further combined with *pAtPGP4::3xVenus-NLS* in the binary vector pAGM4723 (Engler *et al*., 2014).

#### Vectors containing the SBT promoters

The *SBT1.5.2* (2000 bp) and *SBT1.7.3* (1799 bp) promoter regions were each PCR-amplified as three fragments from *P. japonicum* genomic DNA and cloned separately into pAGM1311. The fragments were then combined into the pICH41295 level 0 vector. The *SBT1.1.1* (1691 bp), *SBT1.2.3* (1955 bp), *SBT1.7.2* (1802 bp), and *SBT1.7.1* (1864 bp) promoter regions were each PCR-amplified as one fragment from *P. japonicum* genomic DNA and cloned into the pICH41295 level 0 vector. The promoter sequences were then assembled into level 1 vectors together with the Venus protein module fused with the nuclear localization signal (NLS), and the 3’UTR and terminator module. We also assembled the actin promoter (Wakatake *et al*., 2018) into a level 1 vector together with the mCherry protein module fused with the membrane localization signal (Syntaxin of Plant 122, SYP122), and the 3’UTR and *HSP* terminator module, to generate the *pACT::3xmCherry-SYP122* transcription unit. Each *pSBT::3xVenus-NLS* unit was further combined with *pACT::3xmCherry-SYP122* in the binary vector pAGM4723.

#### Vectors containing Epi10 sequence

A codon-optimized Epi10 construct containing the signal peptide-encoding region of *AtSBT1.7* (At5g67360) (Schardon *et al.*, 2016) was amplified with Golden Gate compatible primers and cloned into pAGM9121 to generate a level 0 CDS1 module (Engler *et al*., 2014). The *Epi10* level 0 CDS1 module was then assembled between the *PjSBT* promoter modules and an *HSP* terminator sequence (Wakatake *et al*., 2018). The final level-1 constructs were combined with the fluorescent transformation marker *p35S:3xVenus-NLS* in the binary vector pAGM4723.

### Transformation of *P. japonicum*

Transformation of *P. japonicum* was performed as previously described by Ishida *et al.* (2011) with several modifications. Silwet L-77 (Bio Medical Science) was added to an *A. rhizogenes* bacterial solution (OD600 = 0.1) to a final concentration of 0.02% (v/v), just prior to transformation. Six-day-old *P. japonicum* seedlings were immersed in the bacterial/Silwet L-77 solution and submitted to ultrasonication using a bath sonicator (Ultrasonic Automatic Washer; AS ONE) for 10 to 15 seconds. The sonicated seedlings were vacuum infiltrated for 5 min. The seedlings were transferred to freshly made co-cultivation medium (Gamborg B5 agar medium with 1% sucrose and 450 µM acetosyringone) and kept in the dark at 22°C for 2 days. After co-cultivation, the seedlings were transferred to B5 agar medium containing cefotaxime (300 µg/mL). After 3 to 4 weeks, the transformed roots were used for infection. Identification of the transgenic roots was performed as previously described by Ishida *et al.* (2016).

### Microscopy

Microscopy with transformed *P. japonicum* was performed as previously described by Spallek *et al.* (2017) with minor modifications. *P. japonicum* with transgenic hairy roots were transferred to water agar plates (0.7% agar; INA) for an additional 2 days before infection of 7-day-old *A. thaliana* seedlings. Infecting plants were then analyzed by confocal microscopy (Leica, TCS SP5 II).

### RT-qPCR

For extraction of total RNA from the haustoria, *P. japonicum* seedlings were grown vertically for 9 days followed by incubation for 2 days on water agar plates before infection of 7-day-old *A. thaliana* seedlings. At 3 and 7 dpi, haustoria were excised and immediately frozen in liquid nitrogen. We removed the *A. thaliana* roots as much as possible. Ten to twenty haustoria were used for each sample. For the 0 dpi samples, we used the root elongation zones from *P. japonicum* seedlings without infection. Total RNA was extracted using the RNeasy plant mini kit (QIAGEN), followed by cDNA synthesis using the ReverTra Ace qPCR RT Kit (TOYOBO). During RNA extraction, we treated with DNase to remove residual genomic DNA. RT-qPCR was performed as previously described by Spallek *et al.* (2017). *PjUBC2* was used as a reference gene. The expression level of each gene was quantified using the ddCt method.

### Primers

All primers used for library preparation, cloning, and RT-qPCR are listed in Supplemental Table S4.

### Statistics

Welch’s *t* test was performed in Microsoft Excel 2016.

### Accession Numbers

Sequence data from this article can be found in the GenBank/EMBL libraries under accession number BankIt2316603: MT149970-MT150066 (SBTs); BankIt2324477: MT226912 (ICSL1), MT226913 (GLP1), MT226914 (CDR1).

MIAME-compliant (minimum information about a microarray experiment) raw RNA-seq data were deposited at the DNA Data Bank of Japan (https://www.ddbj.nig.ac.jp/index-e.html) under accession number SAMD00216292-SAMD00216327.

