## Supplemental file for "Subtilase activity in the intrusive cells mediates haustorium maturation in parasitic plants"

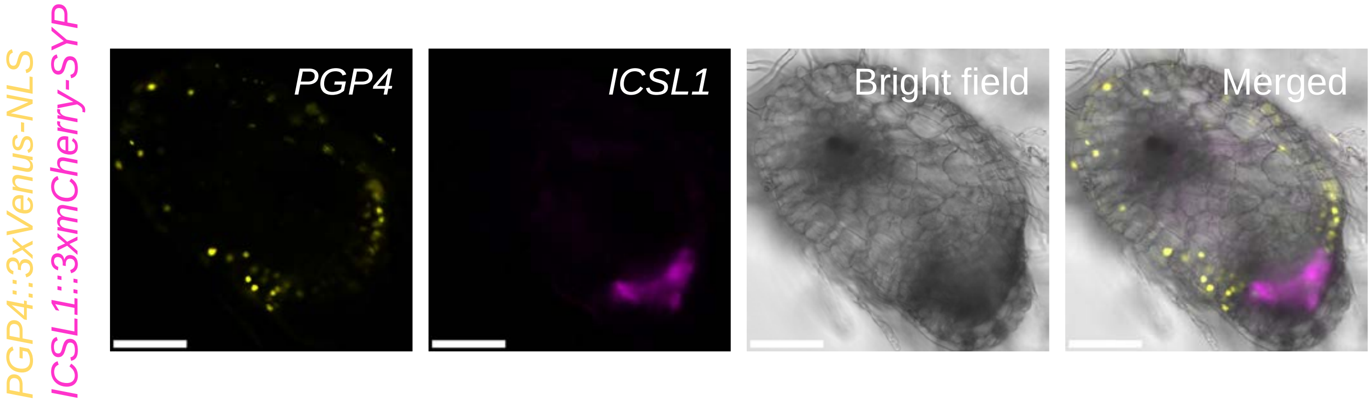


Supplemental Figure S1 **Expression of epidermis and intrusive cell markers during haustorium development.** Expression patterns of the *PGP4* and *ICSL1* promoters in *P. japonicum* during haustorium development at 4 dpi. Bar = 100 µm.

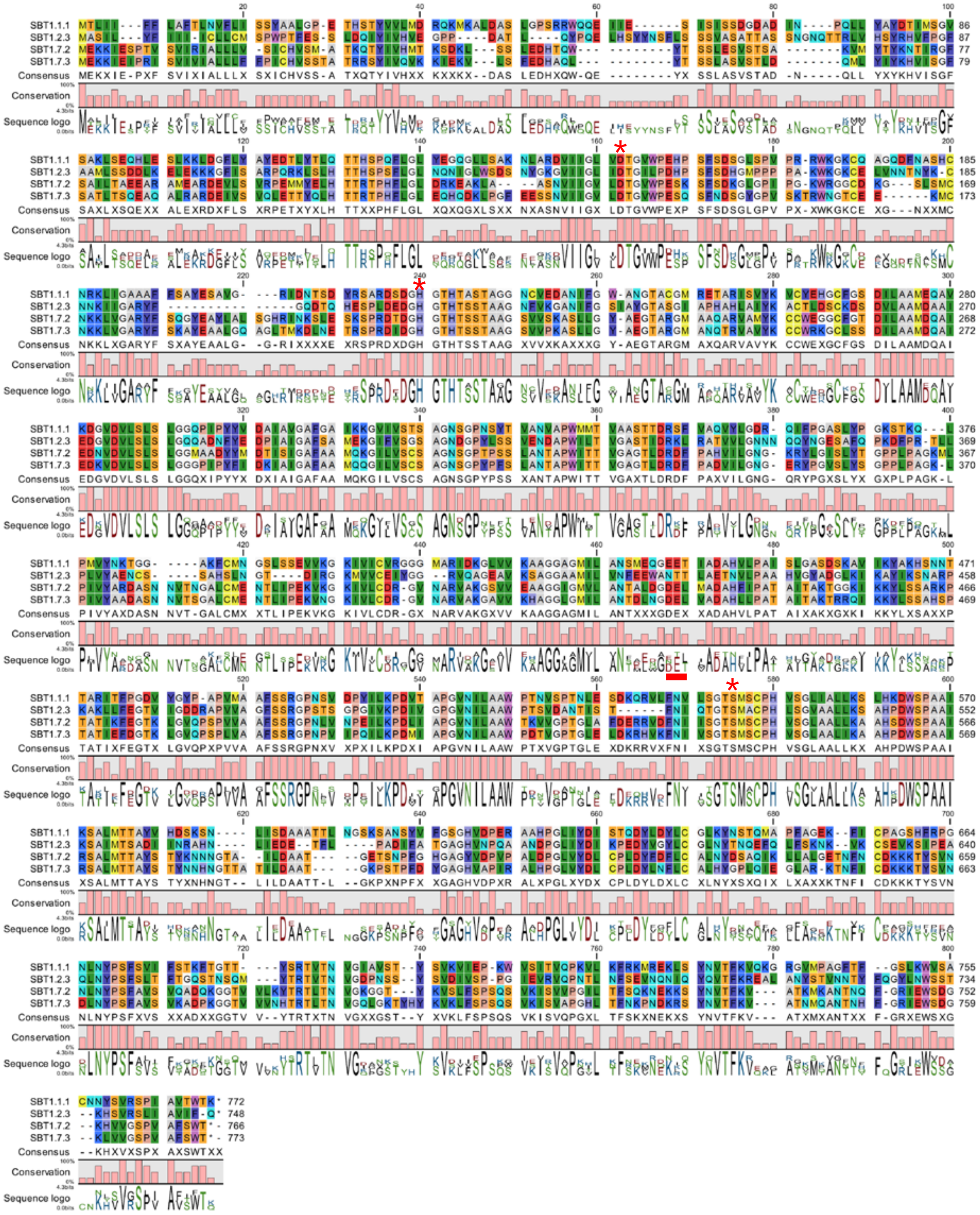


Supplemental Figure S2 **Alignment of amino acid sequences of 4 SBTs whose gene expression was high in intrusive cells.** The asterisks and underline indicate the Asp-His-Ser catalytic triad and the protease domain interface, respectively.

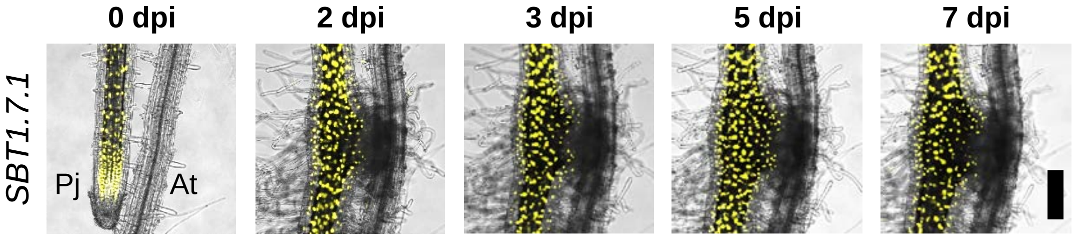


Supplemental Figure S3 **Expression dynamics of the SBT1.7.1 promoter during haustorium development.** Expression patterns of the *SBT1.7.1* promoter driving expression of the Venus fluorescent module in *P. japonicum* during haustorium development at the indicated time points. Bright-field and Venus fluorescent images are merged. Pj, *P. japonicum* root; At, *A. thaliana* root. Bar = 200 µm.

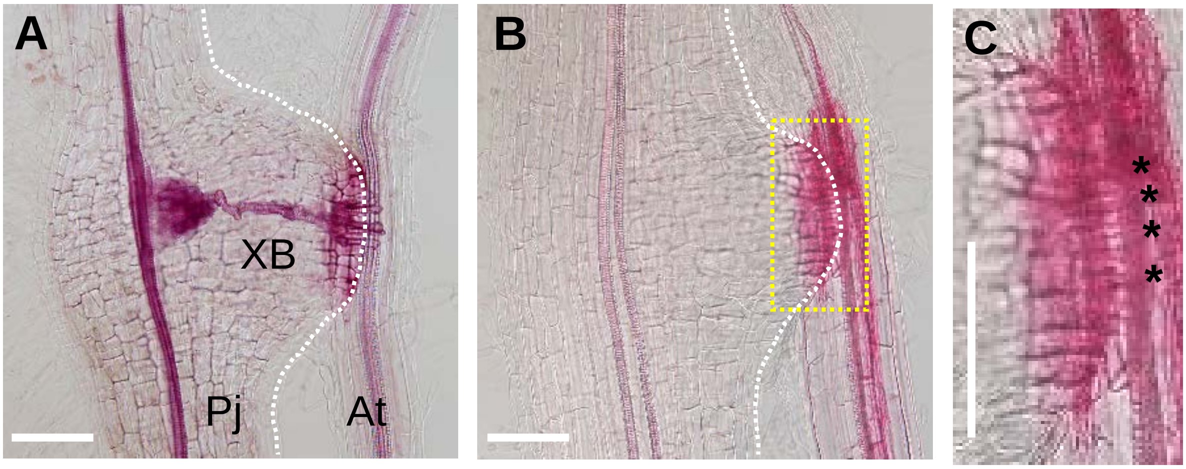


Supplemental Figure S4 **Xylem bridge in the haustoria.** (A) Representative image of a Safranin-O-stained haustoria that that formed a xylem bridge (XB) at 5 dpi. The dashed white line indicates the edge of the haustorium. (B) Representative image of a Safranin-O-stained haustoria that did not form a XB at 5 dpi. (C) Magnified view of the interface region of the haustorium indicated with the dashed yellow line in B. Black asterisks indicate intrusive cells. Pj, *P. japonicum*; At, *A. thaliana*; XB, xylem bridge. Bar = 100 µm.

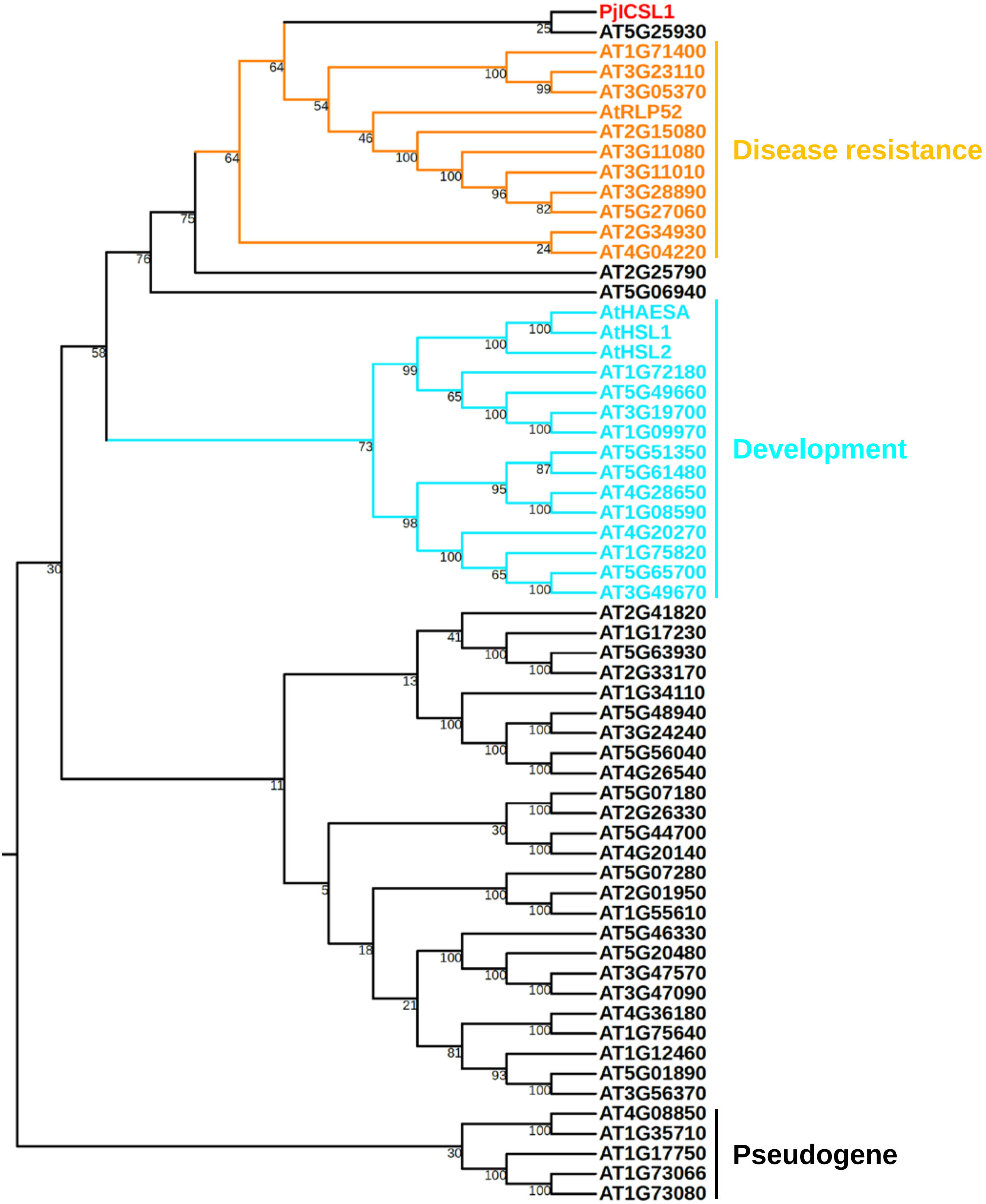


Supplemental Figure S5 **Phylogeny of ICSL1 in *P. japonicum* and the related LRR-RLKs in *Arabidopsis*.** The LRR-RLKs in *Arabidopsis* were selected by protein BLAST analysis against ICSL1 with an E value of < 1E-50. The ICSL1 in *P. japonicum* is shown in red. RLP52 is known to elicit resistance to Powdery Mildew Pathogen (Ramonell *et al*., 2005). HAESA and HSL2 regulate abscission of organs (Stø *et al*., 2015). Putative pseudogenes represent the outgroup. The LRR-RLKs mediating disease resistance and development are shown in orange and blue, respectively.

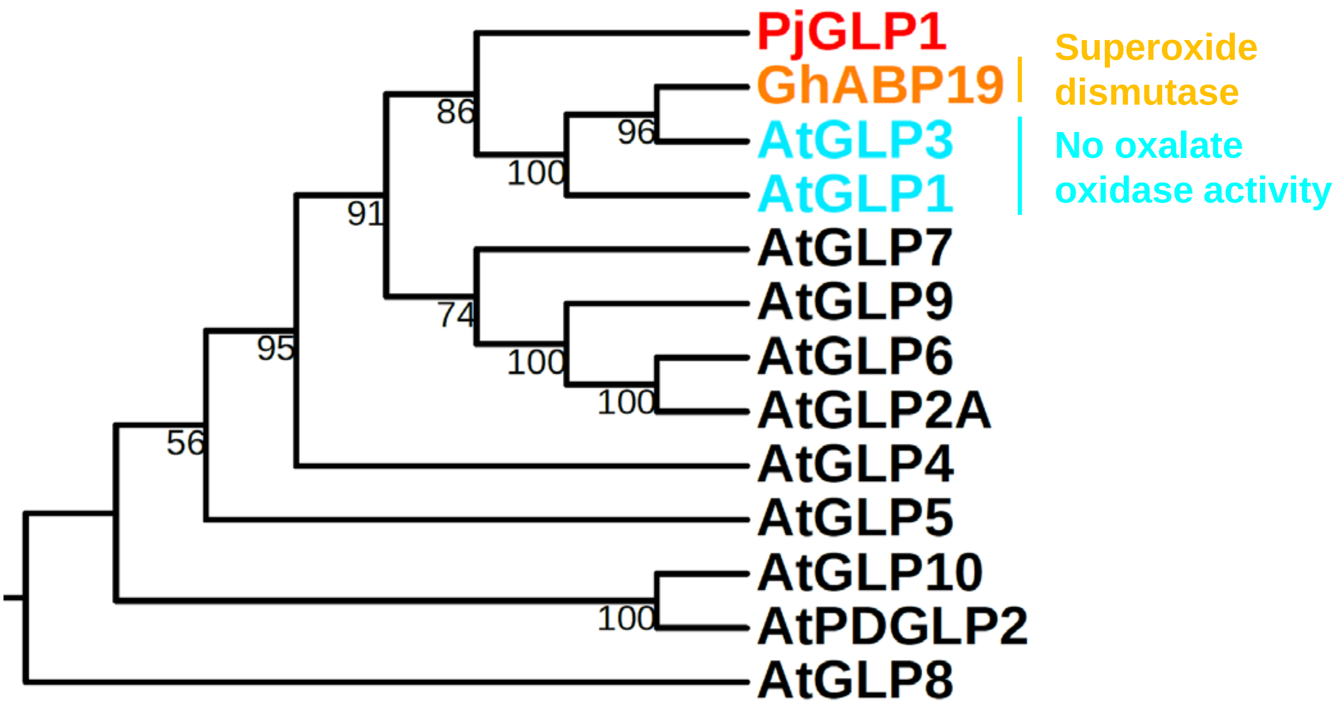


Supplemental Figure S6 **Phylogeny of GLP1 in *P. japonicum*, GhABP19 in *Gossypium hirsutum*, and the GLPs in *Arabidopsis*.** ICSL1 in *P. japonicum* is shown in red. The GLPs with superoxide dismutase activity or without oxalate oxidase activity are shown in orange and blue, respectively.

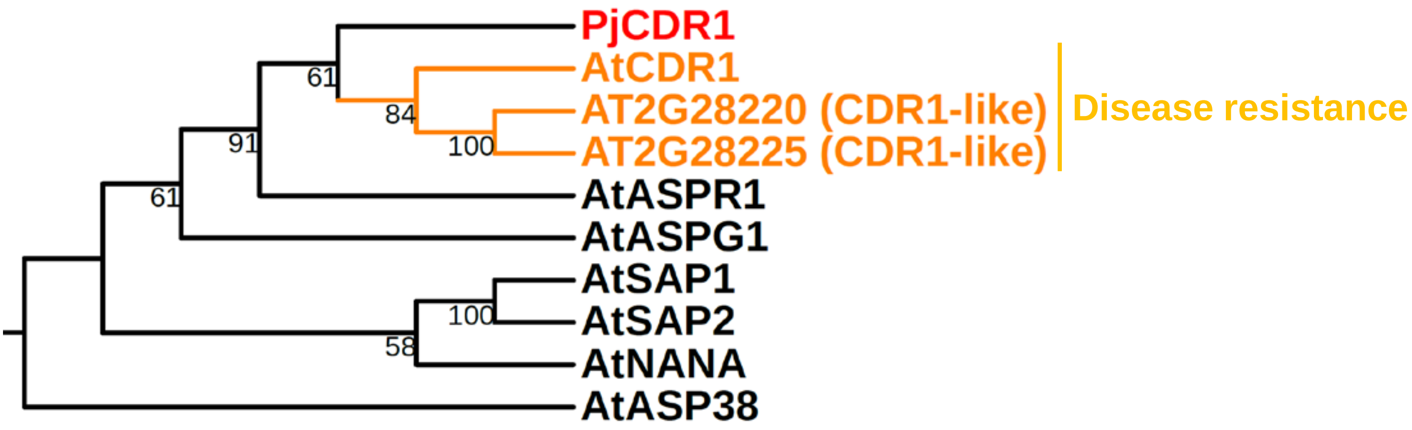


Supplemental Figure S7 **Phylogeny of CDR1 in *P. japonicum* and aspartic proteases in *Arabidopsis*.** CDR1 in *P. japonicum* is shown in red. Aspartic proteases mediating disease resistance and development is shown in orange.

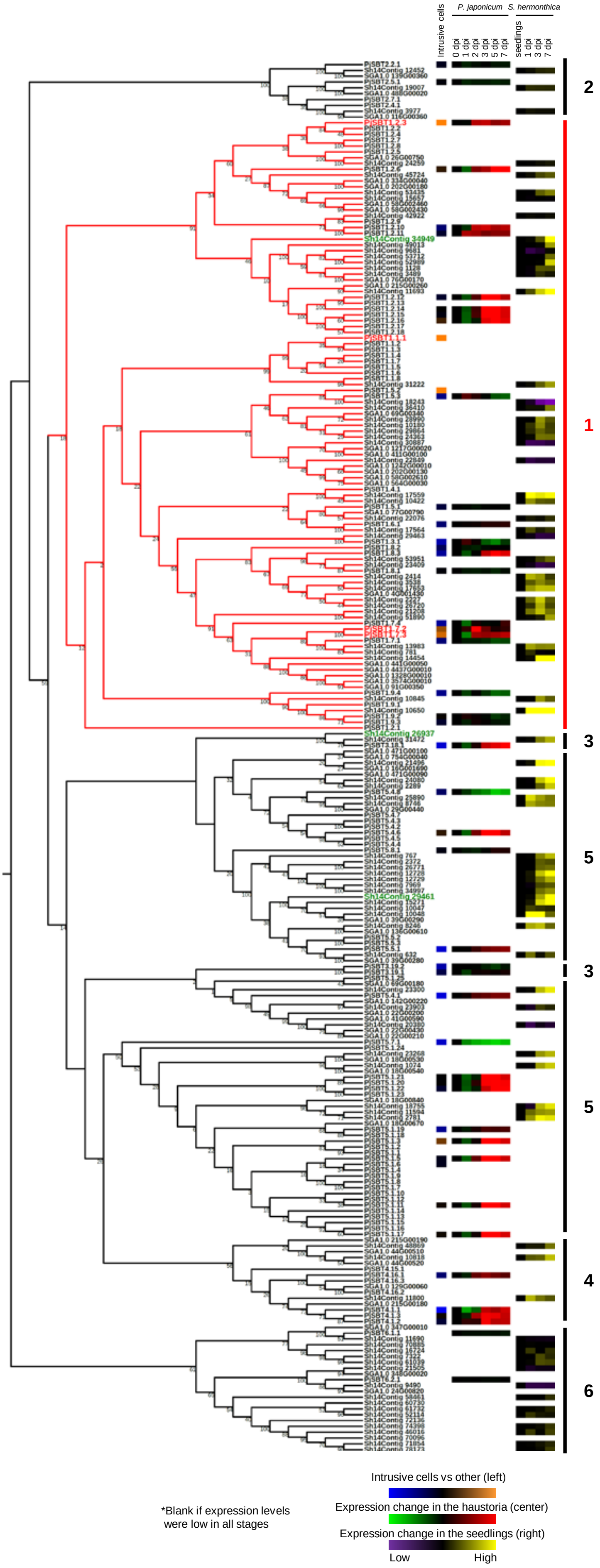


Supplemental Figure S8 **Phylogeny of the SBTs in *P. japonicum*, *S. asiatica* and *S. hermontica*.** The numbers of SBTs in *P. japonicum*, *S. asiatica* and *S. hermonthica* are 97, 50 and 92, respectively. According to Rautengarten *et al*. (2005), SBTs are categorized into 6 groups. The Group 6 SBTs are the outgroup. The tree for the Group 1 SBTs is shown in red. The SBTs expressed in the intrusive cells of *P. japonicum* are shown in red, and the SBTs that are up-regulated at the late stage in *S. hermonthica* are shown in green (Yoshida *et al*., 2019).

The blue/orange squares (left) indicate gene expression ratios of the *SBTs* in intrusive cells relative to the expression in other haustorial parts. The green/red squares (center) indicate gene expression levels of the *SBTs* in the haustoria of *P. japonicum* (0 dpi = 1). The violet/yellow squares (right) indicate gene expression levels of the *SBTs* in seedlings of *S. hermonthica* (seedlings = 1). The transcriptome analysis in *S. hermonthica* was performed by Yoshida *et al*. (2019).

Supplemental Table S1 **List of GO terms enriched in intrusive cells.**

| Go ID | GO term |
|---|---|
| GO:0005618 | cell wall |
| GO:0005886 | plasma membrane |
| GO:0005576 | extracellular region |
| GO:0003674 | molecular function |
| GO:0008152 | metabolic process |
| GO:0019748 | secondary metabolic process |
| GO:0006950 | response to stress |
| GO:0005215 | transporter activity |
| GO:0009607 | response to biotic stimulus |
| GO:0009719 | response to endogenous stimulus |

Supplemental Table S2 **List of GO terms enriched in the haustorial parts, excluding the intrusive cells.**

| Go ID | GO term |
|---|---|
| GO:0005576 | extracellular region |
| GO:0005618 | cell wall |
| GO:0009579 | thylakoid |
| GO:0015979 | photosynthesis |
| GO:0019748 | secondary metabolic process |
| GO:0003824 | catalytic activity |
| GO:0003700 | transcription factor activity, sequence-specific DNA binding |
| GO:0008152 | metabolic process |
| GO:0005975 | carbohydrate metabolic process |
| GO:0003674 | molecular function |
| GO:0009607 | response to biotic stimulus |
| GO:0004872 | receptor activity |
| GO:0016020 | membrane |
| GO:0005886 | plasma membrane |
| GO:0019825 | oxygen binding |
| GO:0006629 | lipid metabolic process |
| GO:0030246 | carbohydrate binding |
| GO:0009719 | response to endogenous stimulus |
| GO:0009628 | response to abiotic stimulus |
| GO:0006091 | generation of precursor metabolites and energy |
| GO:0009058 | biosynthetic process |
| GO:0005215 | transporter activity |
| GO:0009991 | response to extracellular stimulus |
| GO:0007154 | cell communication |

Supplemental Table S3 **Genes exclusively expressed during the parasitic stage.** The transcriptome analysis was performed by Ishida *et al*. (2016). Ten genes that showed the highest expression are listed. Genes encoding SBTs are highlighted in orange.

| Feature ID | $\log_2$ (1+RPKM) | Description |
|---|---|---|
| Pj3contig_3988 | 7.96 | **subtilase family protein** |
| Pj3contig_27632 | 6.91 | ** No description available ** |
| Pj3contig_52149 | 6.74 | **subtilase family protein** |
| Pj3contig_41594 | 6.59 | **subtilase family protein** |
| Pj3contig_3531 | 6.54 | **subtilase family protein** |
| Pj3contig_6540 | 6.02 | disease resistance-responsive protein |
| Pj3contig_49579 | 5.86 | **subtilase family protein** |
| Pj3contig_49870 | 5.74 | ** No description available ** |
| Pj3contig_1736 | 5.60 | **subtilase family protein** |
| Pj3contig_6068 | 5.57 | **subtilase family protein** |

### Supplemental Table S4 **Primers used in this study.**

| Name | Direction | Sequence (5' -> 3') | Purpose |
|---|---|---|---|
| RT primer | | TATAGAATTCGCGGCCGCTCGCGATAATACGACTCACTATAGGGCGTTTTTTTTTTTTTTTTTTTTTTTT | 1st strand synthesis |
| Tagging primer | | TATAGAATTCGCGGCCGCTCGCGATTTTTTTTTTTTTTTTTTTTTTTTT | 2nd strand synthesis |
| PCR primer | | GTATAGAATTCGCGGCCGCTCGCGAT | PCR enrichment |
| ICSL1pro_Lv-1_part1 | Forward | TTGGTCTCAACATggagATCTAATAACCCACGATCGCA | Promoter + 5'UTR |
| | Reverse | TTGGTCTCAACAAcgacGCGTGACAACCGAC | |
| ICSL1pro_Lv-1_part2 | Forward | TTGGTCTCAACATgtcgCTGAAGTCGTGGCAT | Promoter + 5'UTR |
| | Reverse | TTGGTCTCAACAAcattGCGACGGCTCAGGG | |
| GLPpro_Lv-1_part1 | Forward | TTGGTCTCAACATggagTATCACGCTCTCATCTAC | Promoter + 5'UTR |
| | Reverse | TTGGTCTCAACAAacagAAGGAAAAAAAAAGGGGA | |
| GLPpro_Lv-1_part2 | Forward | TTGGTCTCAACATctgtTTTCCTTTTCTCTTTCCG | Promoter + 5'UTR |
| | Reverse | TTGGTCTCAACAAcattAGTCTTTGTTAATGGGTGTG | |
| CDRpro_Lv-1_part1 | Forward | TTGGTCTCAACATggagGGAACCGCCGAGAGATATA | Promoter + 5'UTR |
| | Reverse | TTGGTCTCAACAAaagtTGAGTCCATATGTGAGAG | |
| CDRpro_Lv-1_part2 | Forward | TTGGTCTCAACATacttCTCACAGTCTACTCAAC | Promoter + 5'UTR |
| | Reverse | TTGGTCTCAACAAcattGATGATAAGATTGTATGATACAC | |
| SBT1.1.1pro_Lv0 | Forward | TTGAAGACAAGGAGTATCAGCTAACAATTATCAGGTATCAGCTACC | Promoter + 5'UTR |
| | Reverse | TTGAAGACAACATTTATTGTTGTTGCTGTAAGGAAAAATAAGACGGATG | |
| SBT1.2.3pro_Lv0 | Forward | TTGAAGACAAGGAGAGAAAAAACAAAGTAAAATAGACAAAGGTTCTATCTGCTTTG | Promoter + 5'UTR |
| | Reverse | TTGAAGACAACATTCTGTAACAATCAGACAATGGCTTGCTC | |
| SBT1.5.2pro_Lv-1_part1 | Forward | TTGGTCTCAACATggagAAATAGGCCCAAAACCGACGCC | Promoter + 5'UTR |
| | Reverse | TTGGTCTCAACAAgtatTTTTTACTTATTACTATTTTGTTTCACTTATTACACACTTTTGTTTTGCC | |
| SBT1.5.2pro_Lv-1_part2 | Forward | TTGGTCTCAACATatacATAACAGTATTTGCACATCAAAACAAGAAAC | Promoter + 5'UTR |
| | Reverse | TTGGTCTCAACAAgcgaCCAAAAGAGTCGTAAAGACAG | |
| SBT1.5.2pro_Lv-1_part3 | Forward | TTGGTCTCAACATtcgcTCTCTTTTTTTTTTGTTCGTTTCTTTTTCAGACCCTCTCTATATAATAATATAACCCAGAGCTAATTAAGAAGATCTCATTTCACAAGCACAATTTCATCCTTCCTATATTCGTTTCAGAATGTTGTTGAGACCAA | Promoter + 5'UTR |
| | Reverse | TTGGTCTCAACAAcattCTGAAACGAATATAGGAAGGATGAAATTGTGCTTGTGAAATGAGATCTTC | |
| SBT1.7.2pro_Lv0 | Forward | TTGAAGACAAGGAGAAAAAATATTTTGAAAAATAAATCAGATCAAGCCATTGATTTTAATAAGC | Promoter + 5'UTR |
| | Reverse | TTGAAGACAACATTTTTGTATTTTAGAATTAATTTTGCCAGAGAGGCTG | |
| SBT1.7.3pro_Lv-1_part1 | Forward | TTGGTCTCAACATggagGTTTAAAACGCTACTTATAAGGCGACAC | Promoter + 5'UTR |
| | Reverse | TTGGTCTCAACAAggacAACTTGATAGAAAAATACTTTAGGAGGC | |
| SBT1.7.3pro_Lv-1_part2 | Forward | TTGGTCTCAACATgtccTCTCAAGTATGAGTTTTTTAAAGTTGCCTCC | Promoter + 5'UTR |
| | Reverse | TTGGTCTCAACAAtttgAAACTCAACACCTTTTCAATTTTCTTCACTTG | |
| SBT1.7.3pro_Lv-1_part3 | Forward | TTGGTCTCAACATcaaaGACCACGCTCAGTTGTACACCTC | Promoter + 5'UTR |
| | Reverse | TTGGTCTCAACAAcattCTGGTCCAGGGTGGACACCG | |
| SBT1.7.1pro_Lv0 | Forward | TTGAAGACAAGGAGGACATATTCACCATATGTGCACTTACTTCTGAAACCGAATAAATTTTATAAAG | Promoter + 5'UTR |
| | Reverse | TTGAAGACAACATTCTTTTTTTATTTTCTTAATTTATGCCAGGGATTGTAGAGATGAAAGACTGTGAAG | |
| SBT1.1.1_CDS | Forward | ATGACACTCATCATATTCTTCCTAGCTTTCAC | Coding region |
| | Reverse | TTACTTGGTCCAGGTCACAGCAATAG | |
| SBT1.2.3_CDS | Forward | ATGGCTTCCATATTATATTTTATCATTATCATTTGCCTG | Coding region |
| | Reverse | TCACTGAAATATAACTGCGATAAGGCTC | |
| SBT1.7.2_CDS | Forward | ATGGAGAAGAAAATCGAAAGTCCAACAGTTTCAG | Coding region |
| | Reverse | CTACGTCCAACTAAAAGCAACTGGACTCC | |
| SBT1.7.3_CDS | Forward | ATGGAGAAGAAAATTGAAATTCCAAGAATATCAGTGATTG | Coding region |
| | Reverse | TTACGTCCAACTAAAAGCAACTGGACTC | |
| Epi10_CDS_Lv0 | Forward | TTGAAGACATCTCAAATGTCTAGTAGCTTCCTTAG | Coding region |
| | Reverse | TTGAAGACAACTCGaagcCTAAAGTTTTTGCTGCTGC | |
| PjUBC_qPCR | Forward | ACCATAATTGGACCGCCTGA | RT-qPCR |
| | Reverse | ACAGTTGGGGGACTGTTTGG | |
| SBT1.1.1_qPCR | Forward | ATCCGGGCCTGATTTACGAC | RT-qPCR |
| | Reverse | CACCAGCAAAAGGTGCCATC | |
| SBT1.2.3_qPCR | Forward | TGTCGGGGACCCTAACTCAT | RT-qPCR |
| | Reverse | TTGCTAGGGCCTCTCGTTTG | |
| SBT1.7.2_qPCR | Forward | ATCAAATTGCTTGCGCTGGG | RT-qPCR |
| | Reverse | TTAAGCACCACCGTACCACC | |
| SBT1.7.3_qPCR | Forward | CCCTTTACAGATCGAGGGGC | RT-qPCR |
| | Reverse | CCACCCTTGGGATCAGCTTT | |

Underline: Type IIS enzyme recognition site, Red: overhang of a backbone vector, lowercase: overhang of a gene of interest
